# Perspective taking activates one’s own perceptual object processing in infants and adults

**DOI:** 10.1101/2024.08.06.605501

**Authors:** Anna-Lena Tebbe, Katrin Rothmaler, Moritz Köster, Charlotte Grosse Wiesmann

## Abstract

Perspective taking is central to human cognition and interaction. Preverbal infants already seem to consider the perspective of others, and adults do so continually, in parallel with other cognitively demanding tasks. Yet, the representational format in which others’ perspectives are implemented in the mind and brain remains unclear. We addressed this question by using a neural marker that provides a precise index of visual object processing. We presented adults and 12-14-months-old infants with objects flickering at 4 Hz, evoking phase-locked rhythmic activity at the exact same frequency over visual cortex that is highly specific of visually processing the flickering object. The object then became occluded from the participant’s view. Critically, in one condition another person continued to see the object, whereas in control conditions the observer’s view was blocked or no observer was present. Remarkably, both in adults and infants, the frequency-tagged neural response specific of own visual object processing persisted when their own view was blocked and only the other person could see the object, but not in either control condition. These findings provide direct neural evidence that representing another person’s perspective engages our own perceptual object processing and demonstrate that this representational format is already present in infancy.

**Teaser:** Frequency-tagging shows that objects seen only by others activate one’s own object-specific perception in adults and infants.

## Introduction

As highly social beings, we continually need to keep track of the perspectives of others around us. Communicating with others, interpreting their behavior, and deciding about our own next steps — human thought and interaction is hardly imaginable without perspective taking. Despite its centrality to social cognition, a fundamental question remains unresolved: how is another person’s perspective represented in the mind and brain? At the core of this debate is the question of whether others’ perspectives are represented in a perceptual format or only at later more abstract, inferential processing stages.

Traditionally, the ability to reason about others’ mental states has been assumed to emerge late in the preschool years and to rely on abstract inference-based processes that depend on higher cognitive capacities, such as, language and cognitive control (*1*, *2*). However, this account has been challenged by a growing body of evidence suggesting that, in their behavior, preverbal infants already consider other people’s perspectives, including what others see (*3*, *4*), prefer or intend (*5*), know (*6–8*), and possibly even believe (*2*, *9*, *10*), even when this differs from the infant’s own perspective. At the same time, non-replications and alternative explanations of the infant findings have raised doubts about whether these findings truly demonstrate perspective taking, or whether infants instead rely on simpler strategies such as gaze cueing, enhanced attention or memory for jointly attended events (e.g., (*2*, *10–14*). As a result, a central debate persists: do infants genuinely represent others’ perspectives, and if yes, how could they do so before higher cognitive capacities develop?

One influential proposal is that representing another person’s perspective may involve the activation of perceptual processes used to represent one’s own sensory experience. On this view, tracking what another person sees may reactivate our own object-specific perceptual representations, rather than relying exclusively on abstract, propositional inference (e.g.,*15*–*17*). Representing what another person sees in a perceptual format would make this content available in the same representational space as the infant’s own perceptual experience, providing bottom-up access to this content without explicit, abstract inference.

Support for perceptual accounts has primarily come from behavioral findings that the perspective of others can influence perceptual judgements. For example, adults recognize or detect an object faster when another person is looking at it (*15*, *17*) or falsely believes it to be present (*18*). Similarly, infants’ expectations and processing of their environment are modulated by what others around them see or believe (*18–20*). However, such general modulations cannot determine whether another person’s perspective is represented in a perceptual or inferential format. The presence or gaze of another agent may enhance attention or memory without engaging perceptual representations of the object itself (e.g., *11*, *14*, *20*, *21*).

Neuroscientific research has identified a network of brain regions and temporal dynamics associated with perspective taking and theory of mind, including medial prefrontal and temporoparietal areas (*23*). However, this work has largely focused on where and when perspective-related processing occurs, rather than on how the content of another person’s perspective is represented. In particular, it remains unknown whether representing what another person sees engages one’s own perceptual processing, or whether this information is only available in a higher-order, non-perceptual format.

Here, we directly address this question using a neural marker that provides a precise index of perceptual object processing. We employed a frequency-tagging paradigm in which a visual object flickered at a constant rate (4 Hz), eliciting rhythmic brain activity over visual cortex at the exact same frequency (referred to as steady-state visually evoked potential, SSVEP) (*24–26*). Because this response is phase-locked to the external stimulus and does not occur spontaneously without rhythmic visual input as a trigger (*27*, *28*), it provides a highly specific neural signature of visual processing of the flickering object.

We applied this approach in adults (experiment 1) and in 12- to 14-month-old infants (experiment 2). Participants viewed videos in which an agent observed a flickering object that subsequently became occluded from the participant’s view. In one condition, the agent continued to see the object behind an occluder, whereas in the control condition the object moved into a tunnel that blocked visual access for both the agent and the participant (Fig. 1, Video S1). If representing another person’s visual perspective involves activating one’s own object-specific perceptual processes, then participants should continue to show the specific phase-locked neural response at the object’s flicker frequency after the object disappears from their own view—but only when the agent can still see the object. In contrast, no such response should be present when the agent’s view is also blocked.

**Fig. 1.**
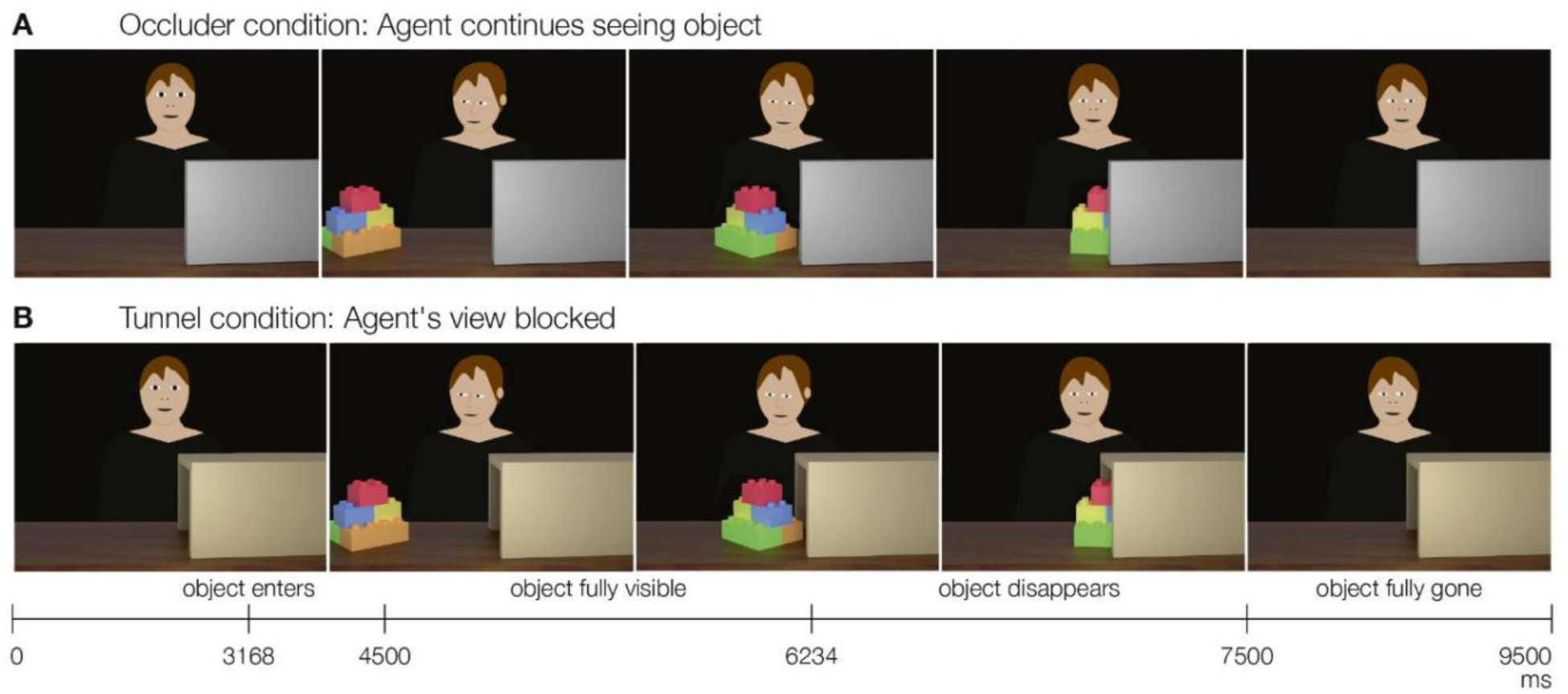
Schematic illustration of the time course of two exemplary trials. (**A**) Flickering objects either moved behind an occluder (continued visual access for the agent only) or (**B**) into a tunnel (neither participant nor agent have visual access).

By testing these predictions, the present study examines whether perspective taking engages one’s own perceptual processing, and whether this representational format is already present in infancy. Demonstrating such an effect would provide direct neural evidence bearing on longstanding debates about the nature of perspective taking and the format in which others’ mental states are represented.

## Results

The objects were flickered 4 times per second. To compare participants’ SSVEP between conditions at every time point during the trial, we therefore preregistered the envelope of their phase-locked 4 Hz activity over the visual cortex using the Hilbert transformation as a measure of the amplitude of their 4 Hz response evoked by the flickering object. This measure of amplitude was used in adults (in experiment 1). In infants (experiment 2), this measure could not be used because of an unbalanced number of trials across the different conditions (see Methods). For the infants, we therefore computed the signal-to-noise ratio (SNR) as a baseline-free measure of their evoked 4 Hz response, which is recommended when the overall power spectrum differs across conditions (*29*) and which we had preregistered as an alternative measure. In order to allow for a direct comparison between infants and adults, we also report the SNR in adults, yielding highly comparable results to the amplitude (reported in the Supplementary Material, S1).

Both adults’ (*N* = 40) and infants’ (*N* = 56) evoked 4 Hz response increased as a function of the object’s visibility to the participant (adults: Fig. 2C; infants: Fig. 3C). This established that viewing the flickering object, moving through the static scene, indeed evoked phase-locked rhythmic activity at 4 Hz, located over occipital sites (Fig. 2B, Fig. 3B). Further, both samples’ frequency spectra (computed during the *object phase*, defined as the time window in which the object was fully visible) showed clear peaks at 4 Hz and its harmonics, as expected for SSVEPs (Fig. 2A, Fig. 3A).

**Fig. 2.**
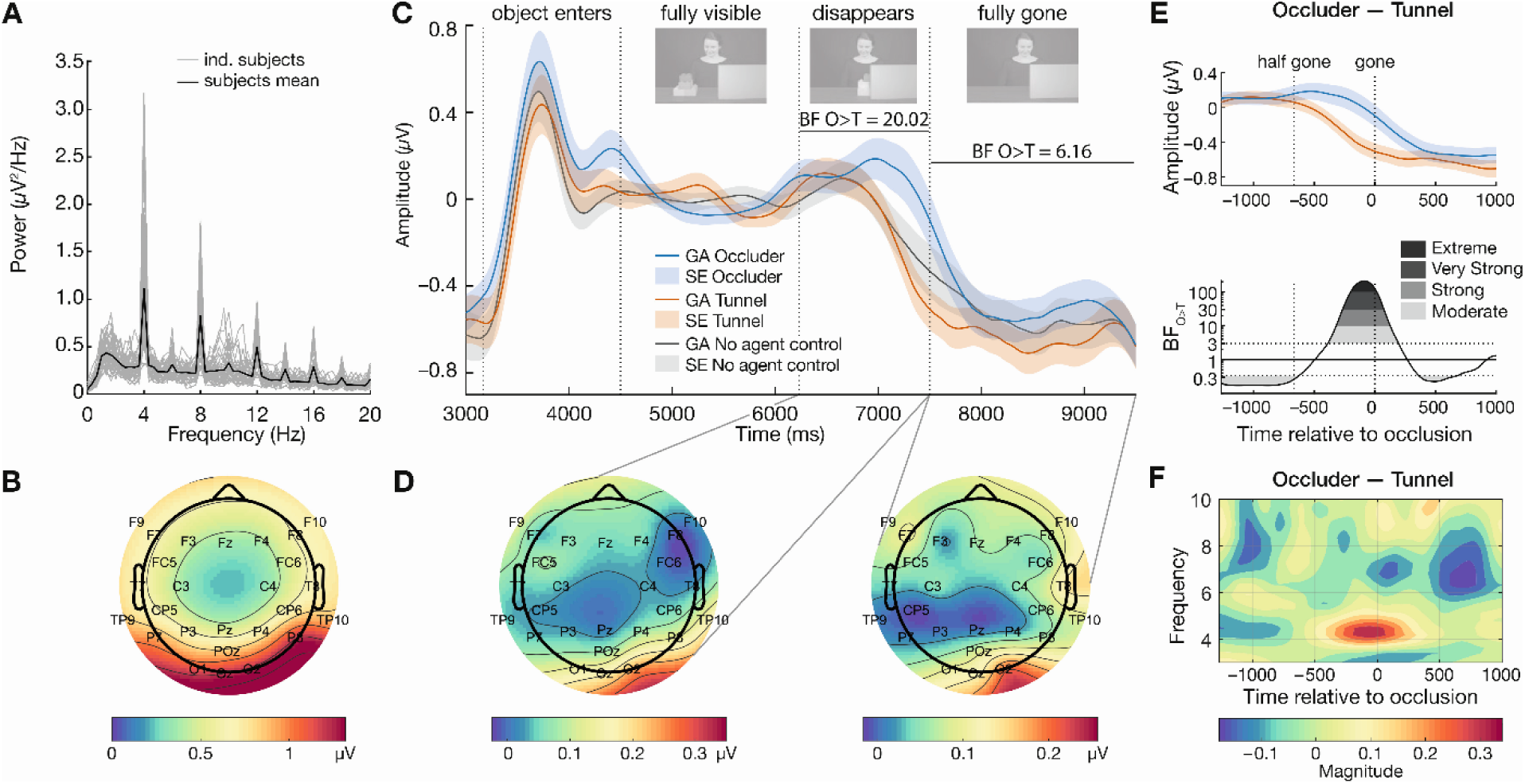
Results of experiment 1: Evoked 4 Hz response in *N* = 40 adults. (**A**) The frequency spectrum shows clear peaks at 4 Hz and its harmonics during the object phase (electrode Oz). (**B**) The topographical results demonstrate a clear evoked 4 Hz response to the flickering object over the visual cortex. (**C**) Baseline-corrected average evoked 4 Hz response envelope over occipital sensors (O1, O2, Oz). Results show a higher evoked 4 Hz response when the agent continued to see the object in the Occluder condition (blue) compared to the Tunnel condition (orange) and a No Agent control (grey), both in the disappearance phase (strong evidence) and during the full occlusion phase (moderate evidence). Shading indicates the standard error. (**D**) Topography of the evoked 4 Hz Occluder-Tunnel difference while the object disappears (left) and during full occlusion (right). (**E**) Evoked 4 Hz amplitude envelope in the period from object disappearance until 1000 ms after full occlusion (top, time relative to full occlusion) and BF_10_ of the dynamic comparison between conditions (bottom) showing higher 4 Hz response in Occluder compared to Tunnel trials (BF_10_ between 3–424.45). Note that the y-axis of the BF_10_ is log-scaled. (**F**) The time-frequency plot displays the power difference between the Occluder and Tunnel conditions at the end of the trial, with a clear 4 Hz peak before and after the object was fully occluded.

**Fig. 3.**
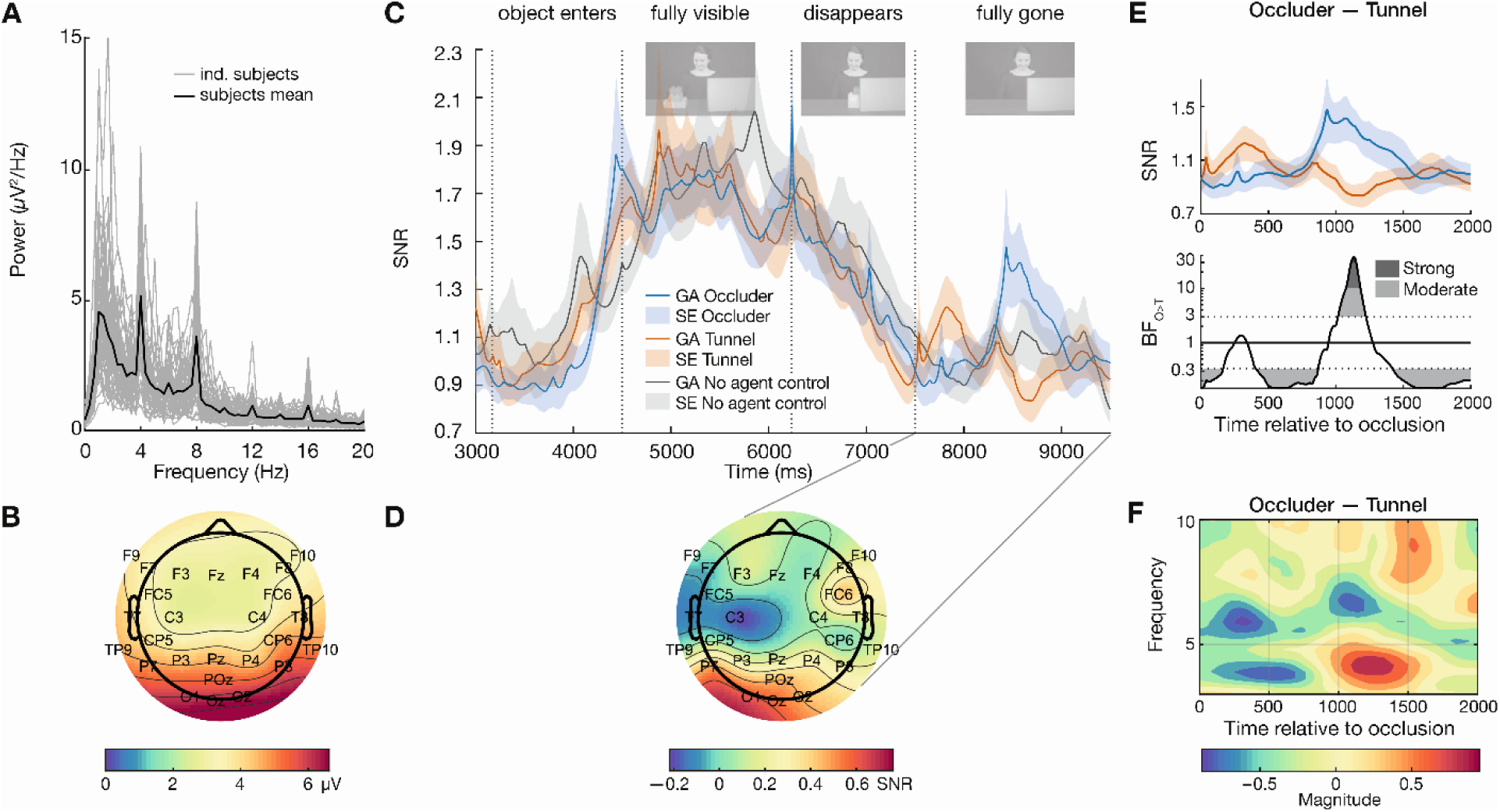
Results of experiment 2: Evoked 4 Hz response in *N* = 56 infants. (**A**) The frequency spectrum shows clear peaks at 4 Hz and its harmonics during the object phase (electrode Oz). (**B**) The topographical results show a clear evoked 4 Hz response to the flickering object over the visual cortex. (**C**) Average 4 Hz SNR over occipital sensors (O1, O2, Oz). Results show a higher evoked 4 Hz response when the agent sees the object in the Occluder condition (blue) compared to the Tunnel condition (orange), but not in a No Agent control (gray), in a time period after full occlusion. Shading indicates the standard error. Note that the short peak around 6200 ms in the Occluder condition is the result of outliers at this specific time point and the difference between condition at this time point is not significant. (**D**) Topography of the evoked 4 Hz Occluder-Tunnel difference during full occlusion. (**E**) Evoked 4 Hz response in the time window from the object’s disappearance until the end of the trial (top, time relative to full occlusion) and the BF_10_ of the dynamic comparison between conditions (bottom). This shows a higher 4 Hz response in Occluder compared to Tunnel trials (BF_10_ between 3-74.45). (**F**) The time-frequency plot displays the power difference between the Occluder and Tunnel conditions at the end of the trial with a clear 4 Hz peak after full occlusion.

### Experiment 1: Others’ perspective engages object-specific perceptual processes in adults

Our central prediction was that this evoked 4 Hz response would also be present after object occlusion when the agent continued to see the object (Occluder condition), as opposed to when her view was blocked by the tunnel (Tunnel condition). Indeed, in adults there was extremely strong evidence for a higher evoked 4 Hz response in the Occluder condition compared to the Tunnel condition, starting during the time window when the object disappeared and lasting until after the object was fully occluded (dynamic Bayesian t-tests: BF_10_ ranging between 3 and 424.45, from 7050 ms to 7736 ms, see Fig. 2C & 2E). These effects remained when the evoked 4 Hz response was averaged within two time intervals: while the object was fully occluded (*full occlusion phase*, defined as the first frame in which the object was fully occluded until the end of the trial: Bayesian Wilcoxon signed-rank test: BF_10_ = 6.16, W = 568, *R̂* = 1.011), and while the object was disappearing (*disappearance phase*, defined as a time window of 1333 ms (80 frames) starting with the first frame in which the object began to disappear: BF_10_ = 20.02, W = 601, *R̂* = 1.023). Similar analyses using the SNR instead of the envelope (for comparison with the infants) confirmed the results obtained with the envelope, with strong evidence for a higher 4 Hz SNR in the Occluder compared to the Tunnel condition in a similar time window (see S1).

Adults’ visually evoked response to the flickering object was thus prolonged when the agent continued to see the object and remained present throughout the occlusion phase, despite the fact that the object was no longer visible to the participants themselves. This indicates that adults processed what the other person saw similar to seeing the flickering object themselves.

### Experiment 2: Others’ perspective reactivates object-specific perceptual processes in infants

If perspective taking activates our own perceptual processes, this perceptual format may also already provide infants with bottom-up access to the content viewed by others, despite their still immature cognitive capacities. In experiment 2, we therefore presented 12- to 14-month-old infants with the same videos and compared their evoked 4 Hz response in the Occluder versus the Tunnel condition. As in adults, this yielded very strong evidence for higher evoked 4 Hz responses in the Occluder compared to the Tunnel condition. In infants, this occurred in a time window after the object was fully occluded (dynamic Bayesian t-tests: BF_10_ ranging from 3 to 74.45 between 8460 ms and 8752 ms, Fig. 3). Averaged across the entire disappearance and full occlusion phase, no evidence for a difference was found (full occlusion phase: Bayesian Wilcoxon signed-rank BF_10_ = .66, W = 629, *R̂* = 1.005; object disappearance phase: BF_10_ = .18, W = 509, *R̂* = 1.000). This suggests that, for infants, the effect was specific to the observed time window after full occlusion.

Thus, similar to adults, infants showed a 4 Hz response evoked by the flickering object, even though the object was no longer visible to them, but only remained visible to the other person. However, infants showed this effect only after the object was already fully occluded. In adults, in turn, the effect was already present while the object was in the process of disappearing, in form of a prolongation of their own visual response to the object, and then lasted throughout the object occlusion (see Fig. S2 for a comparison of effect sizes).

### Excluding differential eye-movements

#### Frontal electrodes

To exclude that the observed difference between the Tunnel and Occluder conditions may have resulted from differential eye movements between the conditions, we analyzed frontal electrodes as these capture eye movements. Over frontal electrodes, no difference between conditions was found, neither in adults nor in infants, including when eye movements were retained in the signal (see S3). This confirms that the observed effects did not result from differential eye movements between the conditions, which would have been visible in frontal electrodes.

### Excluding differences in spontaneous theta-band activity

Beyond differential eye-movements, attending to different contents could, in principle, engage spontaneous, intrinsic theta-band processes that may have inflated the evoked 4 Hz response. However, because any such spontaneous processes would not be phase-locked to the object, their signatures should emerge in the *induced* (i.e., non-phase-locked) rather than the evoked response (*30*). To exclude this, we therefore additionally examined the induced 4 Hz activity (see S4). This showed no increase either in the induced 4-Hz rhythm nor in the broad-band theta power (see S5), confirming that the observed effects indeed reflect differences in the object-specific perceptual SSVEP response.

### No Agent control condition

Although the visual features of the occluder and tunnel (e.g., surface area, luminance) were closely matched and their colors counterbalanced, the minimal visual differences required to show an occluder versus a tunnel could have contributed to the difference in the evoked 4 Hz signal, in particular, during the disappearance phase. To rule out this possibility, we additionally acquired a non-social control condition recorded in the same adult and infant participants. In the non-social control, the object moved behind the occluder, but no agent was present watching the object (see Video S3). When no agent was present, the evoked 4 Hz response was neither prolonged nor enhanced compared to the tunnel condition (see Fig. 2 and 3). Specifically, both in adults and infants, there was moderate evidence for the absence of a difference between the Tunnel and the No Agent control condition (adults: dynamic t-tests BF_10_ = .17–.62 between 7050-7736 ms; Bayesian Wilcoxon signed-rank test for the disappearance phase: BF_10_ = .19, W = 440, *R̂* = 1.000; and the full occlusion phase: BF_10_ = .34, W = 344, *R̂* = 1.002; infants: dynamic t-tests: BF_10_ = .15–.71 between 8460-8752 ms; full occlusion phase BF_10_ = .17, W = 398, *R̂* = 1.001; disappearance phase: BF_10_ = .70, W = 291, *R̂* = 1.002). In contrast, there was moderate evidence for a greater response in the Occluder compared to the No Agent control condition during the disappearance phase in adults (dynamic t-tests BF_10_ = 3–19.99 between 6958-7450 ms; disappearance phase: BF_10_ = 7.82, W = 570, *R̂* = 1.017; full occlusion phase: BF_10_ = .09, W = 344, *R̂* = 1.000) and inconsistent evidence in infants (disappearance phase: BF_10_ = .10, W = 356, *R̂* = 1.000; full occlusion phase: BF_10_ = .51, W = 491, *R̂* = 1.002. This supports that the extended evoked 4 Hz response to the object in the Occluder compared to the Tunnel condition was not due to visual differences between the tunnel and the occluder but indeed resulted from the agent’s visual access to the object.

### Eye-tracking

While adults showed an immediate prolongation of their evoked 4 Hz response, infants reactivated their perceptual processing of the object only after a delay. To follow-up on potential reasons for the observed difference in timing between adults and infants, we ran additional eye-tracking experiments with independent samples of *N* = 24 adults (experiment 3) and *N* = 22 infants (experiment 4). Participants were presented with the same stimuli as in experiment 1 and 2 while their gaze was recorded with an eye-tracker (details see *Methods*).

To analyze participants’ looking behavior, we defined three areas of interest (AOIs) around (1) where the object was last visible before disappearing (object area), (2) the face area, and (3) the Occluder/Tunnel area (see Fig. 4). Looking behavior was quantified via relative looking scores (RLS) between two AOIs respectively ranging from 0 to 1 (details see *Methods*), with 1 indicating looking only to the first area and 0 only to the second area.

**Fig. 4.**
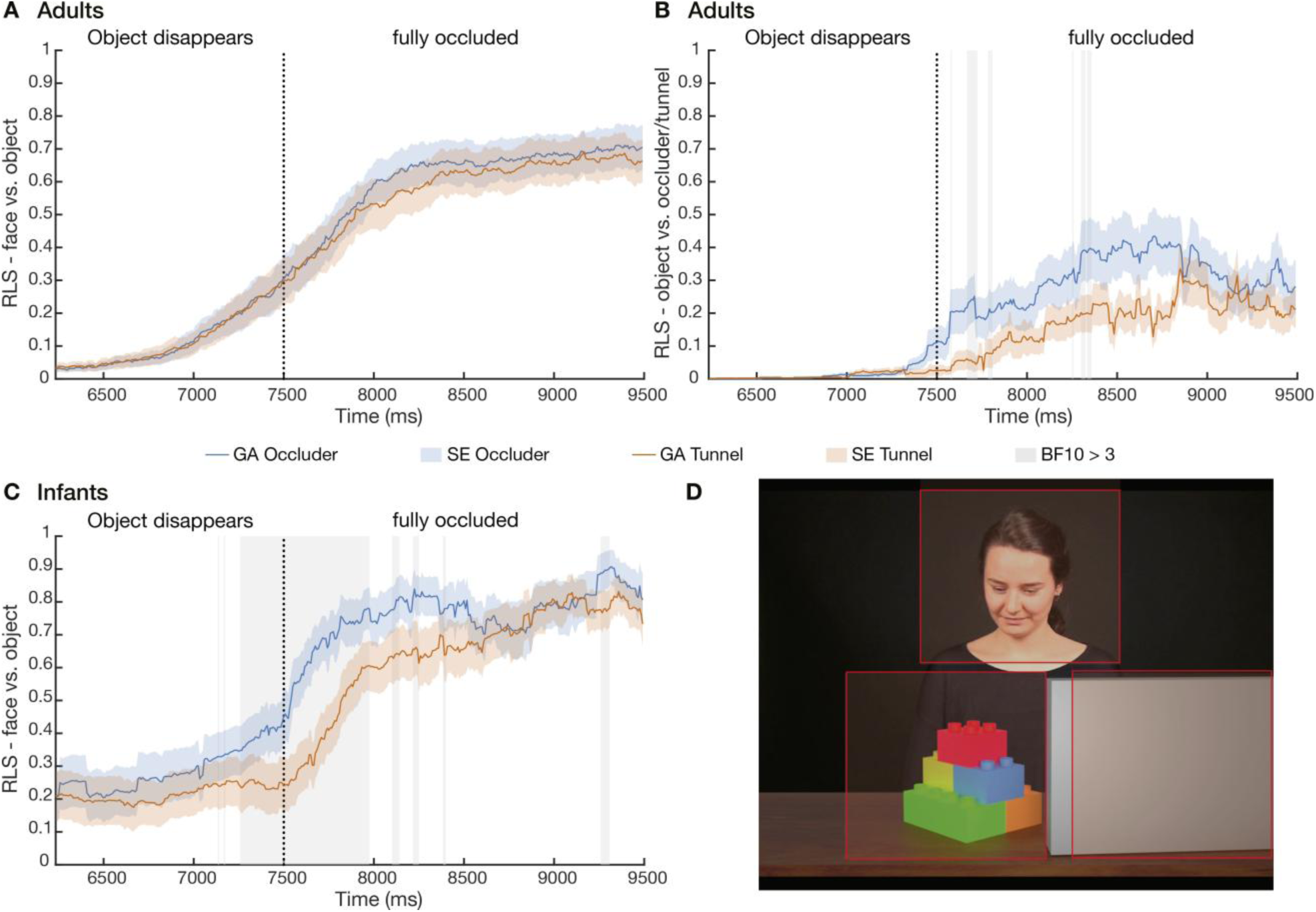
Results of Experiment 3 and 4: Eye-tracking. Relative looking scores (RLS) to different areas of interest (AOI) in the disappearance and full occlusion phase in adults and infants. (**A**) Mean RLS to the face versus object area by condition in *N* = 24 adults, showing no difference between conditions. (**B**) Mean RLS to the object versus occluder/tunnel area in adults. On average in the full occlusion phase, adults looked relatively more to the occluder in the occluder compared to the tunnel condition (BF_10_ = 4.24). Gray shadings indicate timepoints where the Bayesian t-tests showed evidence for a condition difference in dynamic t-tests (BF_10_ between 3 and 8.8). (**C**) Mean RLS to the face versus object area for infants. On average, infants showed higher relative looking to the face area in the occluder compared to the tunnel condition, both while the object disappeared (BF_10_ = 4.29) and after its full occlusion (BF_10_ = 6.31, dynamic t-tests: gray shadings indicate BF_10_ between 3 and 768.99). Note that only very few infants looked into the occluder/tunnel area at all (*N* = 6 or 7 depending on condition). (**D**) AOIs for Experiment 3 and 4 (face area, object area, occluder/tunnel area).

### Experiment 3: Adult eye tracking

Adults showed no condition differences in relative looking to the face area compared to the object area (disappearance phase: BF_10_ = 1.18; full occlusion: BF_10_ = .49) or the occluder/tunnel area (disappearance phase: BF_10_ = .55; full occlusion: BF_10_ = .09, details see S7). However, in the full occlusion phase, participants showed higher relative looking to the occluder area than the object area in the occluder condition compared to the tunnel condition (disappearance phase: BF_10_ = .24; full occlusion phase: BF_10_ = 4.24; mean RLS_obj>occl_ = .68, *SD* = .33; Tunnel: mean RLS_obj>occl_ = .83, *SD* = .18; dynamic t-tests BF_10_ = 3–8.80 between 7583 - 8358 ms, see Fig. 4). This indicates that adults followed the trajectory of the occluded object more throughout its occlusion when the agent could still see it.

### Experiment 4: Infant eye-tracking

Infants, in contrast, showed higher relative looking to the face area compared to the object area in the occluder compared to the tunnel condition, both while the object disappeared (BF_10_ = 4.29; W = 130; *R̂* = 1.001; Occluder: mean RLS_face>obj_ = .30, *SD* = .36; Tunnel: mean RLS_face>obj_ = .21, *SD* = .33) and after its full occlusion (BF_10_ = 6.31; W = 161; *R̂* =1.002; Occluder: mean RLS_face>obj_ = .75, *SD* = .25; Tunnel: mean RLS_face>obj_ = .65, *SD* = .26). This was the case from shortly before the object had fully disappeared approximately until the time point when infants reactivated their object-specific SSVEP in experiment 2 (dynamic t-tests: BF_10_ = 3-768.99 between 7266 – 7975 ms and repeatedly between 8108 – 8400 ms, see Fig. 4). In contrast to adults, infants did not show any difference in relative looking to the object area compared to the occluder/tunnel area (details see S7).

This indicates that, when the agent continued to see the object, infants first focused on the agent and only after this, reactivated their own neural visual processing of the object. Adults, in contrast, followed the trajectory of the object more throughout its occlusion when the agent could still see it, in line with the immediate and continuous prolongation of their evoked 4 Hz response. Importantly, these differences in gaze strategy cannot explain the condition differences in the evoked 4 Hz response. As shown in the control analyses above, we ruled out that condition effects in the EEG stemmed from differential eye movements (as these would have been visible in frontal electrodes) and found evidence against differences in the non–phase-locked theta activity, excluding an alternative explanation in terms of spontaneous theta-band responses driven by differences in attended stimuli.

## Discussion

Our findings demonstrate that, when observing another agent seeing an object, both adults and infants show a neural signature that is highly specific of their own visual processing of the object, even though their own view of the object is blocked. Specifically, flickering stimuli are known to elicit synchronized rhythmic activity over visual cortex at the exact same frequency and phase of the stimulus, referred to as SSVEP (*24*, *25*). In contrast to spontaneous neural oscillatory activity, SSVEPs are synchronous in phase and phase-locked to the stimulus (*24*, *25*, *31*). They therefore provide a highly specific neural signature that can be directly linked to perceiving the flickering stimulus (*24*). The phase-locked 4 Hz response in our study therefore directly reflects participants’ perceptual processing of the flickering object. Both adults and infants continued to show this highly specific *perceptual* response to the object when another agent continued to see it, although participants no longer saw the object themselves. This was not the case when the agent’s view was also blocked, or no agent was present. This provides direct neural evidence that we process what only others can see by activating our own object-specific visual processes of it.

Importantly, alternative explanations according to which following another person’s gaze merely enhances attention to the object in form of a directional cue (*14*, *32*, *33*) are ruled out by our design. The object was no longer visible to the participant, precluding enhanced attention to the object. Moreover, the agent’s gaze direction was identical between both experimental conditions, and the agent attended the object equally before its occlusion. This also speaks against a recent account that, when infants attend to an object together with an agent, this leads to better encoding of the object while it is visible (*11*). Instead, in our paradigm, infants need to infer the agent’s visual access, and our findings show that they then actually *represent* what the other sees. However, they do so by reactivating their own visual representation of the object.

An additional No Agent control condition confirmed that the effect did not arise from visual differences between the occluder and tunnel. Indeed, when no agent was present, the evoked 4 Hz response was neither prolonged nor enhanced after the object had moved behind the occluder. Together, these controls demonstrate that the observed neural response reflects the agent’s visual access to the object rather than low-level attentional or perceptual differences.

By showing that perspective taking activates our own perceptual processing of the content seen by another person, our findings address a longstanding debate concerning the representational format of others’ mental states. Rather than relying exclusively on higher-order inferential processes, others’ perspectives are represented in a perceptual format, making their content directly available to perception-based cognitive processes, such as judgments and action selection. Perception-based actions and decisions are known to be faster and more robust under cognitive load than those relying on higher-order reasoning (*34*). This is consistent with the faster feed-forward dynamics of perceptual neural pathways compared with higher-order association systems (*35*). Thus, representing others’ visual content in a perceptual format may enable rapid, efficient behavioral responses. This is in line with behavioral findings that adults consider others’ perspectives quickly and in parallel to other cognitively demanding tasks, such as, while speaking (*36*) or acting (*37*, *38*). Moreover, representing others’ visual perspective in a perceptual format may provide infants with a bottom-up route to accessing what others see, circumventing the need for higher-order cognitive processes. While our findings provide direct neural evidence for perceptual representations during perspective taking, future work will need to establish their causal role in perspective taking, for example, by perturbing perceptual processing using brain stimulation methods.

If we process what others see in the same format as our own perception, the question arises how the brain differentiates this from seeing it ourselves. In adults, the evoked 4 Hz response to the object was considerably lower when only the agent saw the object than while it was still visible for the participant. This suggests differences in signal strengths between the representation of others’ perspectives and perception, as recently also observed for imagination (*39*). Further, the perspective of others may only be encoded at specific frequencies (see S4-S6). In addition, identifying our perceptual representation of the object *as the agents’ perspective*, distinct from our own, likely requires additional neural mechanisms. Potentially these involve higher-order association regions typically implicated in mentalizing, such as the default network (*23*). Recent evidence suggests that mentalizing engages visual brain regions prior to higher-order areas (*40*), and it has been proposed that the default network may decouple representations from perception (*41*). This raises the possibility that perceptual representations provide an initial format that may subsequently be integrated with higher-level processes.

Indeed, infants may represent what others see perceptually, without necessarily distinguishing this from their own view. Consistent with this possibility, infants have been shown to display altercentric memory errors in their behavior, misremembering object locations based on where another person last saw them rather than where the objects actually were (*42*, *43*). Thus, perceptual representations may provide infants with a short cut to accessing the content of others’ perspective despite their still immature higher cognitive capacities. Indeed, there is research indicating that perspective taking processes in infants may be different from those that support explicitly reasoning about others’ mental states as distinct from our own (*44*, *45*). In adults, the perceptual format may likewise offer a fast route for accessing others’ perspectives without explicitly representing them as distinct from one’s own, explaining why others’ perspectives can interfere with one’s own perception and modulate behavior even later in development (*15*, *17*, *18*, *46*, *47*). Importantly, previously used behavioral and neural indicators (*15*, *18–20*, *42*) were not specific to perception, leaving open the question of the representational format that allows for such modulations.

Adults showed an immediate extension of their perceptual processing of the object when the agent continued to see it. In contrast, infants appeared to reactivate their perceptual processing of the object only after a delay. An additional eye-tracking experiment indicates that, in our paradigm, after object occlusion, infants first directed their gaze to the agent and did so more when she could still see the object than when her view was also blocked. This suggests that infants registered the higher relevance of the agent in the Occluder condition already at object occlusion, but may have needed more time to reactivated their own visual representation of the flickering object, triggering perceptual processing despite fully occluded from their view. Adults, in turn, followed the occluded object more when the agent could still see it, in line with their continuous perceptual processing of the object throughout its disappearance and occlusion. An exciting avenue for future research is to investigate the developmental trajectory of these changes from infancy to adulthood. In addition, future work combining EEG and eye tracking simultaneously within the same individuals could directly test how variability in looking behavior relates to the timing and strength of perceptual object representations.

As the perceptual representation is maintained, despite not seeing the object oneself, it is likely also available for other mental states than the visual perspective. For example, our visual working memory of an object in a certain location may be enhanced in the same way, if an agent has seen the object in a certain location in the past and thus believes it to be there. This would explain why modulations of adults’ and infants’ behavior on the environment have also been found when agents had a false belief (*18*, *19*, *37*, *42*).

Young infants have immature higher cognitive capacities but are strongly dependent on others to navigate and learn from a complex environment. Having access to what others attend to, or hold in mind, even when infants cannot see it themselves, is essential for predicting caregivers’ actions, following what they attend, and understanding what they refer to in communication. Processing others’ perspectives perceptually, within one’s own representation of the world, may provide a mechanism for infants to understand others and learn from them without the cognitive demands of maintaining multiple distinct representations. And not only infants. Our findings suggest that this mechanism persists into adulthood, offering a feed-forward route for considering others’ perspectives when cognitive resources are limited. Such intuitive alignment with others from early on in life may be a driver of human cooperation and at the basis of our unique capacity to learn from and teach others (*48*, *49*). As such, the revealed mechanism may constitute an adaptive advantage leading to the ultra-social nature of human cognition, forming a foundation for the evolution of human culture (*48*, *50*).

## Materials and Methods

The video stimuli, experimental procedure and data processing was identical in Experiment 1 (adults) and Experiment 2 (infants) unless otherwise specified.

### Experiment 1 (adults)

#### Sampling approach

Our sample size was based on a preregistered Bayesian sequential testing scheme (*51–54*). In Bayesian sequential testing, researchers collect data until a predefined minimum sample size of participants is reached. If the Bayes factor meets or exceeds a predefined threshold of evidence for either hypothesis, data collection is stopped (see *50*, *51*). We continued data collection until the Bayes factor (BF) of our main hypothesis (enhanced SSVEP during the object disappearance) reached 3 or 1/3. We prespecified a minimum sample size of N = 30 (*55*) and continued testing until sufficient statistical evidence for or against a difference between the Occluder and Tunnel condition was reached, yielding a sample size of *N* = 31. Because of the complex nature of the preprocessing and analyses and the scheduling procedures, nine additional adult participants had been collected by the time we realized that our stopping criterion had already been reached. We here report the data of the full sample of *N* = 40 participants and report the data at the stopping criterion (*N* = 31) in the SI, yielding comparable results (see S8 and Fig. S6).

#### Participants

Experiment 1 reports data of *N* = 40 healthy adult participants (15 female, 25 male; *Median_age_* = 26.05 years, *SD* = 3.66, also see Table S3). Two additional participants were excluded from the final sample due to poor EEG data quality (more than 50% EEG trials with artifacts). None of the participants reported any history of neurological or psychiatric disorders or photic epilepsy. All participants had normal or corrected to normal vision, provided written informed consent prior to participation and were compensated financially for their time. The study received approval from the Ethics Committee of the University of Leipzig.

#### Stimuli and Procedure

In a within-subject design, we presented participants with videos, in which an agent observed a flickering object moving across a table. Towards the end of each trial, the object disappeared behind an occluder, or in a control condition into a tunnel. Critically, in the Occluder condition, the agent continued to see the object, whereas in the Tunnel condition, her view was blocked. Each trial started with an attention getter presented for 1000 ms (animal picture and sound), followed by a black screen, presented for a random duration between 500 and 800 ms. Subsequently, one of the videos with the flickering objects was shown. The videos featured an actress positioned at the center of the scene. The agent smiled briefly, then gazed towards the left bottom of the table, where an object appeared shortly after. The object moved across the table, followed by the gaze of the agent and was fully visible for 1733 ms. The object then took 1266 ms to move behind the Occluder or inside the Tunnel at the right end of the table. After this, the object was completely occluded for 2000 ms. The duration of the videos was 9.5 sec. In an additional control condition, the object disappeared behind the occluder, but no agent was present watching it.

Participants were presented with 60 videos per condition, organized in blocks of 30 videos of the same condition. The order of the blocks was counterbalanced across participants. Trials featured 20 different objects that were shown in random order. To make the conditions easily distinguishable, Occluder and Tunnel were shown in two different colors (brown and gray) that were matched for luminance and counterbalanced across participants. In order to stress the different spatial properties of Occluder versus Tunnel, and to ensure that participants understood the agent’s visible access to the object depending on condition, each of the blocks was preceded by a familiarization video. These showed how the Occluder or Tunnel (depending on condition), entered the scene, made an 180° turn, and then moved into its final position, while the agent followed it with their gaze.

#### No Agent control

To control for signal differences that may have resulted from the visual difference between the tunnel and occluder rather than the agent’s visual access to the object in the Occluder condition, the same participants were also presented an additional No Agent control condition. This control condition was identical to the Occluder condition, except that a blurred light was shown instead of an agent that matched the luminance of the agent conditions. Further, the videos were 1000 ms shorter as the part featuring the smiling agent before the object entered the scene in the beginning of the other conditions was left out. As for the other conditions, before showing the no agent videos, a familiarization video showed the same occluder movement as the Occluder familiarization video but without an agent.

Videos were created using the graphics software Blender (v. 2.8, (*56*)). The stimuli were presented using Psychtoolbox (version: 3.0.17; (*57*, *58*) in Matlab (R2017a; Mathworks, Inc., Natick, MA, USA) and were played with 60 frames per second (i.e., 16.67 ms refresh interval) at the center of a 17-inch CRT screen at a visual angle of 12° x 22°. To establish a 4 Hz flicker, we presented the frames of our videos at a duty cycle of 7:8 (i.e., seven frames (116.69.ms) showing a bright object (‘on’), followed by eight frames showing the same object strongly dimmed (‘off’), followed by 8 on-frames and 7 off-frames. Importantly, not the whole scene but only the object was presented in a “flickering” mode. Participants viewed the display at a distance of 110 cm and were instructed to watch the videos attentively and to avoid eye movement and blinks during video presentation. To maintain participants’ attention throughout the experiment, they were asked control questions on properties of the object (e.g., color, shape, appearance) after randomly selected trials (two questions within every 10 trials). The questions could be answered with ‘yes’ or ‘no’ by pressing one of two mouse buttons.

#### EEG Recordings and Data Preprocessing

EEG was recorded continuously using 28 active electrodes (FC5/6, Cz, C3/4, T7/8, CP5/6, TP9/10, Pz, P3/4, P7/8, Poz, Oz, O1/2, ground electrode located at FP1, placed according to the International 10–20 System and secured in an elastic electrode cap (Easycap GmbH, Herrsching, Germany). Electrode impedances were kept below 30 kΩ. Data were digitalized with a sampling rate of 500 Hz with Cz as reference and processed offline using Brain Vision Analyzer (Version 2.1.2.327) and MATLAB (2023a; Mathworks, Inc., Natick, MA, USA). We excluded epochs with major artifacts by visual inspection. We further excluded segments with significant artifacts identified via automatic artefact correction (1) gradient (maximal allowed voltage step): 50 µV/ms; 2) difference in amplitude (maximum-minimum): 300 µV in an interval of 300 ms; 3) minimum/maximum allowed amplitude of ±500 µV; 4) lowest allowed activity in intervals of 100 ms: 0.5 µV). Data was band-pass filtered to 1-20 Hz using an IIR Butterworth filter with zero phase shift. Eye Movements were identified and corrected automatically using the Ocular Correction Independent Component Analysis (ICA) integrated in the Brain Vision Analyzer Software and discarded after visual inspection. EEG epochs were segmented from 0 to 9500 ms post-onset.

#### Steady State Visually Evoked Potential Analysis

Artifact-free epochs were averaged separately for each condition. Data were bandpass filtered with a 10^th^ order zero phase Butterworth filter having a halfwidth of 1 Hz around the stimulation frequency. To avoid filter edge artifacts, value padding was performed at the beginning and the end of the signals (500 values on each side). The time-varying amplitude of the SSVEP response was extracted via the Hilbert transform. For the statistical analysis, a cluster average was computed for the region of interest (O1, O2, Oz) based on the expected topography and main cortical source of the SSVEP signal (*59*, *60*). To eliminate offset differences between conditions, the mean amplitude of the object phase (i.e., the time window during which the object was completely visible) was subtracted as baseline.

#### Statistical analyses

Statistical analyses were carried out with Matlab (R2023a; (*61*) and JASP (*62*)). We performed Bayesian analyses and used default priors for all models, i.e., an equal, uninformed prior probability as we had no a priori information regarding the effects. Following conventions, we considered a BF_10_ between 3 and 10 as *moderate* evidence, between 10 and 30 as *strong* evidence, between 30 and 100 as *very strong* evidence, and larger 100 as *extreme* evidence for the given hypothesis H1 (*63*). Analogously, a BF_10_ between 1/3 and 1/10 was considered *moderate* evidence, between 1/10 and 1/30 as *strong* evidence, between 1/30 and 1/100 as *very strong* evidence, and smaller than 1/100 as *extreme* evidence in favor of H0. BFs between 1/3 and 3 were considered inconclusive.

The signal envelopes were analyzed dynamically by calculating Bayesian directed paired t-tests between the conditions at each time point from when the object started disappearing until the end of the trial (*53*, *54*). In addition, the signal envelopes were averaged across two time windows: 1) the *object disappearance phase*, starting from the first time point at which the objects start to disappear until 80 frames [∼1333 ms] later, and 2) the *occlusion phase*, starting from the time point at which the object is completely occluded until the end of the trial. As q-q-plots (see S9) indicated minor deviations from normality, we used we used non-parametric statistical test alternatives whenever indicated and available.

### Experiment 2 (Infants)

#### Participants

Similar to the adults, our final infant data set resulted from a Bayesian sequential testing scheme with a minimum N of 40 and stop criterion of BF = 3 or 1/3 , leading to a sample of *N* = 56 healthy full-term infant participants aged between 12 and 15 months (31 female; *Median_age_* = 13.4 months, *SD* = 22.6 days). Fifteen additional subjects were excluded from the sample because they did not provide at least five artifact-free trials per condition. Due to complex data preprocessing and analyses including manual coding of infant’s looking behavior, which influenced the final sample size as an exclusion criterion, we inadvertently recorded an additional 16 infants after the stopping criterium had been met. In the results, we report the full sample (*N* = 56), but the results at the stopping criterion (*N* = 40) are comparable (as reported in S7). Parents gave written informed consent before participation, and infants received a small gift. The study was approved by the Ethics Committee of the University of Leipzig.

#### Stimuli and Procedure

Infants were presented with the same videos as adults (see Experiment 1). Infants saw a total of up to 40 videos per condition, presented in blocks of 20 videos of the same condition, in counterbalanced order across participants. On average, infants contributed 11.4 trials (± 6.1) in the Occluder condition and 12 trials (± 8.2) in the Tunnel condition. We monitored infants’ attention online via camera. Each block started with the familiarization video that was played repeatedly until the infant had fully seen it once. Infants watched the videos on their parents’ lap with a viewing distance of approximately 80 cm from the monitor.

#### Gaze coding

The experimental sessions were video-recorded for offline coding of infants’ attention to the screen. Thirty percent of the videos were coded by a second independent rater. In case of disagreement, a third rater coded the data, and the rating of the majority was used. The interrater agreement between the first and second ratings was substantial (Median Kappa = .64). Since the SSVEP response reflects a visual signal elicited by on–off stimulation, it is essential to confirm that infants were attending to the screen while the flickering object and its disappearance were visible in order to accurately assess the decay in signal strength. We therefore only included trials in which infants looked at the screen for at least 3 out of 4.3 sec while the object was visible (period from first time visible until occluded). Note that the reported effect remained similar when varying the inclusion criteria (see S10).

#### EEG Recordings and SSVEP Analysis

We recorded a 28-channel EEG with the identical layout used for the adults. There was no difference in the number of trials between the Occluder and Tunnel condition (BF_10_ = .172, error estimation = .072%). Because of strong baseline differences between conditions in the infant sample and an unbalanced number of trials between the social and No Agent control conditions (BF_10_ = 1713.86, error estimation = 3.29×10-10%), we used the signal-to-noise ratio (SNR) as a baseline-free measure of the SSVEP response that it is recommended when the overall power spectrum differs across conditions (*29*). To extract the SNR, the trial averages were bandpass filtered using 10^th^ order zero-phase shift IIR Butterworth filters with a halfwidth of 1 Hz on the frequency of interest (4 Hz) and on the neighboring frequencies (2 and 6 Hz). The SNR was calculated as the ratio between the signal envelope at 4 Hz divided by the average signal envelope at neighboring frequencies (2 and 6 Hz).

### Experiment 3 & 4 (Eye-tracking)

To further investigate the differences in the timing of the evoked 4 Hz effect in adults and infants, we conducted two eye-tracking experiments, one with adults and one with infants, where we showed the same stimuli as in Experiment 1 and 2 but recorded their gaze with an eye-tracker instead of recording EEG.

#### Participants

For Experiment 3, we tested an additional sample with a total of *N* = 22 infants (11 female, 11 male, mean age *M* = 13.5 months, *SD* = 41 days) that did not participate in the EEG experiment. Another nine infants were tested but excluded due to because their gaze could not be tracked (*N* = 7) or because they did not contribute any trials (*N* = 2). For Experiment 4, we recorded an independent sample of *N* = 24 adults (13 female, 9 male, 2 diverse; mean age *M* = 26.21, *SD* = 4.9) that did not previously participate in the EEG experiment. One additional adult participant was recorded but excluded because their gaze could not be tracked.

#### Procedure

Participants’ gaze data was recorded using a Tobii X120 eye tracker (Tobii Technology, Stockholm, Sweden) with 120 Hz sampling rate. The eye-tracker was placed 60 cm away from the participant at an angle of 30°. Before the experimental recording, each participant passed a five-point calibration procedure. We presented participants with the identical videos from the main experiment. Each trial started with an attention getter, followed by a black screen presented for 500 ms before one of the videos was shown. Infants were presented a total of up to 40 videos per condition, presented in blocks of 20 videos of the same condition, adults were presented with 30 videos per condition. Two orders were implemented, so that the first block started with Occluder trials and was followed by the Tunnel condition or vice versa. Before each block, participants were shown two familiarization videos per condition.

We defined three equal-sized areas of interest (AOIs, each (500×460 pixels): (1) the object area (where the object was visible before disappearing), (2) the face area (centered around the agent’s face), and (3) the Occluder/Tunnel area (see Fig. 4). A relative looking score (RLS) between two AOIs was quantified as the looking duration to area A divided by the sum of the looking durations to area A and B (*64*). The proportion of fixations was analyzed in a time window starting from 4500 ms until the end of the trial, corresponding to the beginning of the object phase until the end of the occlusion phase. An average RLS was computed per condition and participant.

Manuscript preparation. ChatGPT version 5.2 was asked to check the Abstract, Introduction and Discussion of the original manuscript concerning grammar, language, style, and flow. The output was checked thoroughly, and some recommendations were followed, while others were discarded.

## Acknowledgments

We would like to thank the infant, parents, and adult participants who volunteered to participate in this study. Further, we would like to thank Ruth Faßbender, Marie Michael, Valentina Tast, Noa Wallraff, Laura Walz, and Zeynep Yenen for their support in data collection and video coding the data. Finally, we would like to thank Frederik Bartholin, Radek Cichy, Joshua Grant, Stefanie Höhl, Andreas Keil, Vadim Nikulin, and Jonathan Phillips for very helpful comments to previous versions of this manuscript.

## Author contributions

Conceptualization: AT, KR, MK, CGW

Methodology: AT, KR, MK, CGW

Software: AT, KR, MK

Validation: AT, KR

Formal analysis: AT, KR

Investigation: AT, KR

Resources: AT, KR, CGW

Data curation: AT, KR

Writing – original draft: AT, CGW

Writing – review & editing: AT, KR, MK, CGW

Visualization: AT, KR, CGW

Supervision: KR, CGW

Project administration: AT, CGW

Funding acquisition: AT, CGW

## Data, code, and materials availability

All data needed to evaluate the conclusions in the paper are present in the paper and/or the Supplementary Materials. Aggregated data and the code are available on OSF (https://osf.io/nakd7/overview?view_only=82759583fac44260b7197cac2c5ee074). Interested researchers may request access to the complete stimulus material on the institute’s servers by contacting the corresponding author.

