## SupplementalMaterial for "Perspective taking activates one’s own perceptual object processing in infants and adults"

Anna-Lena Tebbe *et al.*

**This PDF file includes:**

Supplementary Text
Figs. S1 to S8
Tables S1 to S4
Movies S1 to S5
References (# to #) (if applicable—these should refer only to references in the SM)

### Supplementary Text

#### *S1. SNR analysis - Adults*

For direct comparison with the infant data, we also computed the SNR for the adult sample. This confirmed the results obtained with the envelope, with a higher 4 Hz SNR in the Occluder compared to the Tunnel condition in a similar time window as for the envelope ($BF_{10}$ ranging between 3 and 37.24, from 7146 ms to 7414 ms, see Fig. S1).

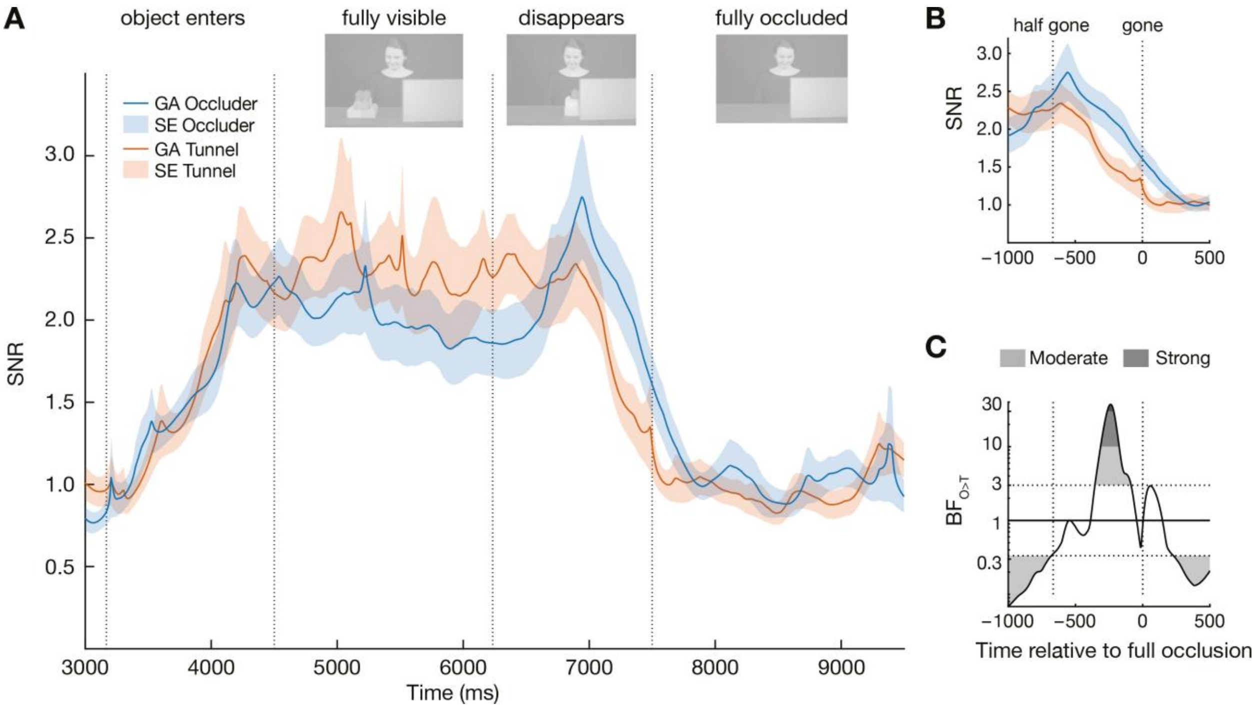


**Fig. S1. Signal-to-noise ratio (SNR) for the adult sample.** (**A**) The SNR of the evoked 4 Hz signal envelope by condition for $N = 40$ adults, averaged over occipital electrodes (Oz, O1, O2). Bayesian t-tests showed moderate to strong evidence for a higher response in Occluder compared to Tunnel trials in the time window from 7146-7414 ms. (**B**) SNR in the time window from 1000 ms before until 500 ms after object occlusion (time relative to full occlusion) and (**C**) the respective $BF_{10}$ of the t-tests comparing Occluder and Tunnel trials ($BF_{10}$ between 3 and 37.24).

*S2. Effect plots*

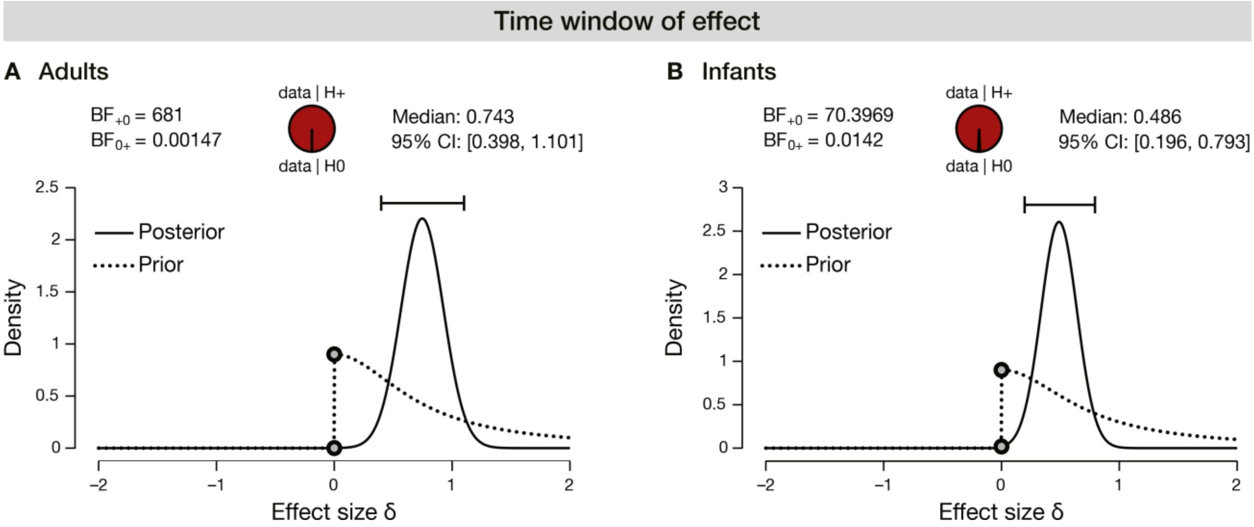


**Fig. S2. Effect size plots for Experiment 1 and 2.** Effect size plots for $N = 40$ adults (**A**) and $N = 56$ infants (**B**), showing a large effect for adults and a moderate effect for infants.

*S3. Frontal electrodes*

To exclude that the observed effect of visual access on the 4 Hz response resulted from differences in eye movements, we conducted the same analyses for a frontal electrode cluster (Fz, F3, F4) as differential eye movements would be reflected in differential signal at frontal electrodes. As we integrated an ICA to correct for eye movement artifacts in our preprocessing, this might have factored out effects of eye movement. We therefore compared the signal at frontal electrodes between conditions without correcting for eye movement artefacts. This revealed evidence against a condition difference over frontal electrodes in adults (disappearance phase: $BF_{10} = .20$, W = 418, $\hat{R} = 1.001$; Occluder: $M = .07$, $SD = .22$; Tunnel: $M = .06$, $SD = .26$; full occlusion phase: ($BF_{10} = .56$, W = 469, $\hat{R} = 1.001$; Occluder: $M = .05$, $SD = .23$; Tunnel: $M = -.01$, $SD = .18$). Similarly, in infants there was evidence against condition differences in the disappearance phase: ($BF_{10} = .09$, W = 688, $\hat{R} = 1.001$; Occluder: $M = 1.15$, $SD = .52$; Tunnel: $M = 1.22$, $SD = .56$) and in the full occlusion phase ($BF_{10} = .06$, W = 600, $\hat{R} = 1.001$; Occluder: $M = 1.00$, $SD = .37$; Tunnel: $M = 1.13$, $SD = .43$). This confirms the occipital focus of the observed effect as expected for SSVEPs and supports that the effect did not result from differential eye movements between conditions.

*S4. Induced activity*

Although we used the SSVEP to obtain a highly specific 4-Hz response reflecting the processing of the object, the SSVEP amplitude could, in principle, be inflated by spontaneous theta activity (e.g., due to differences in looking behavior across stimuli). Because such differences would appear in the induced, non–phase-locked signal, we conducted an additional control analysis of the induced activity in the infant sample to rule out this possibility. Specifically, we explored the stimulus-related changes in the SNR of 4 Hz *induced* activity in contrast to the *evoked* 4 Hz response that is phase-locked to the stimulus onset. To this end, we corrected the envelope of the trial-wise 4 Hz activity by subtracting the trial-average (ERP) before calculating the condition average. This revealed no difference between the Occluder and Tunnel condition, neither during the disappearance phase ($BF_{10} = .188$, $W = 476$, $\hat{\boldsymbol{R}} = 1.001$) nor during the full occlusion phase, ($BF_{10} = .173$, $W = 447$, $\hat{\boldsymbol{R}} = 1.007$), also see Fig. S3. This shows that the observed effect is a stimulus-driven signal phase-locked to 4 Hz, reflecting precisely the rhythmic signal used to tag the object.

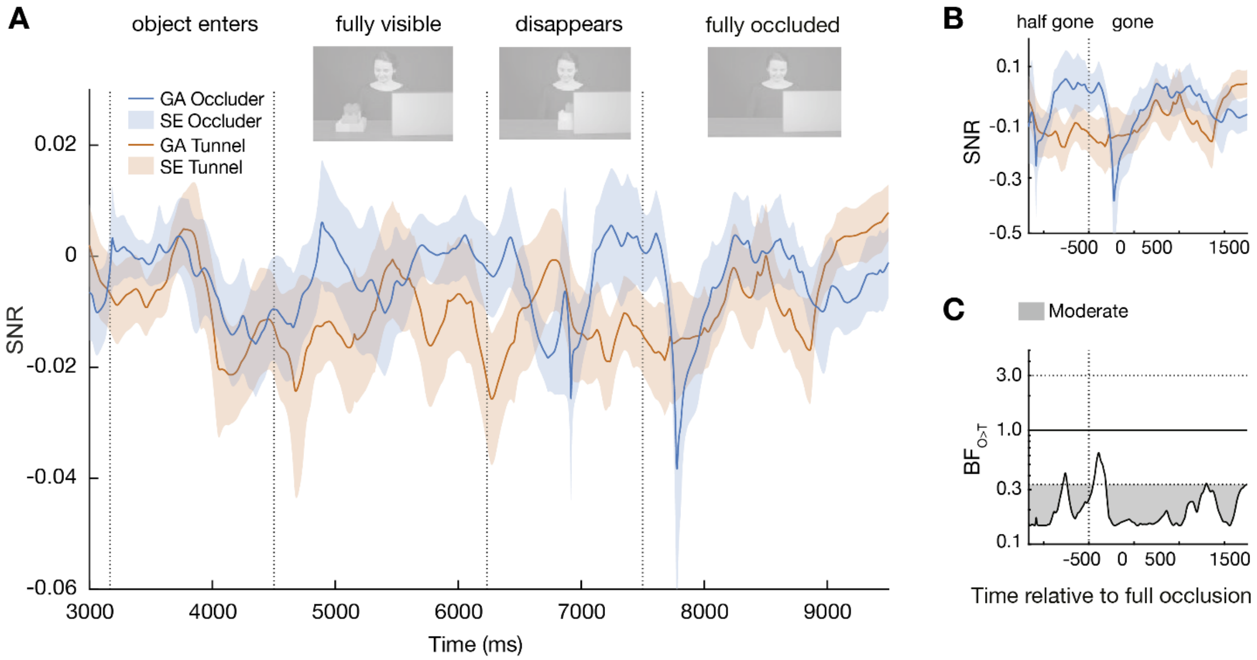


**Fig. S3. Induced 4 Hz rhythm in the infant sample.** (**A**) The SNR of the 4 Hz rhythm after subtracting the ERP on the single-trial level by condition for $N = 56$ infants, averaged over occipital electrodes (Oz, O1, O2). Bayesian t-tests showed no evidence for a difference between conditions, neither during the time window of the evoked effect between 8460 ms and 8752 nor during the rest of the trial ($BF_{10}$ between .15 and 1.34).

*S5. Overall theta band activity*

To confirm the finding that the visual processing of the agent's perspective was specific to 4 Hz and not present in the overall theta activity, we tested for differences between conditions in the overall theta band (i.e., 4 - 8 Hz). To this end, we bandpass filtered the data with a $10^{th}$ order zero phase Butterworth filter having a halfwidth of 3 Hz around the target frequency of 6 Hz and extracted the time-varying amplitude between 4 and 8 Hz via the Hilbert transform. Wilcoxon signed-rank test contrasting the theta band activity in Occluder vs Tunnel trials showed no difference in the full occlusion phase ($BF_{10}$ = .31 W = 429, $\hat{R}$ = 1.000; Occluder: *M* = -.54, *SD* = .52; Tunnel: *M* = -.58, *SD* = .52) or the disappearance phase in adults ($BF_{10}$ = 1.99, W = 541, $\hat{R}$ = 1.002; Occluder: *M* = .04, *SD* = .36; Tunnel: *M* = -.08, *SD* = .37). This supports that the extended evoked 4 Hz response to the object in the Occluder compared to the Tunnel condition was not due to an increase in overall theta band activity in the Occluder condition but was specific to the 4 Hz stimulation frequency. This analysis is not meaningful in infants as we had to use the Signal-to-Noise Ratio (SNR) around the target frequency of 4 Hz and surrounding frequencies as noise as the main measure of the evoked 4 Hz response in infants (see Methods section).

*S6. Results with 6 Hz control frequency*

To test whether the observed effect was frequency specific, we presented the same adult sample with an additional control condition, featuring the same videos but using 6 Hz as alternative stimulation frequency in the same frequency band. We reasoned that what others see may only be processed like perception at specific frequencies that resonate with ongoing internal neural rhythms sustaining relevant cognitive functions (*13–15*). We had chosen 4 Hz as stimulation frequency based on its role in visual working memory (*16–18*), which we reasoned to be important for sustaining the perceptual processing of the object after its occlusion when the agent continued to see it. Indeed, the extension of participants' visual processing of the object as a function of the agent's visual access was specific to 4 Hz and not present at 6 Hz or in the overall theta activity (see also S5). Specifically, while participants showed a clear evoked 6 Hz signal while the objects was visible to them (see Fig. S4A), these evoked oscillations were not prolonged or enhanced after the object's occlusion when the agent continued to see the object (dynamic t-tests: $BF_{10} < .30$, Bayesian directed Wilcoxon signed-rank test for the disappearance phase: $BF_{10} = .11$, $W = 359$, $\hat{R} = 1.001$; full occlusion phase: $BF_{10} = .10$, $W = 350$, $\hat{R} = 1.000$; see Fig. S4C). Thus, processing the agent's perspective like perception was specific to 4 Hz, a frequency that as a spontaneous rhythm in the brain supports visual working memory. This suggests that the external stimulation at 4 Hz resonated with participants' ongoing 4 Hz rhythm, supporting sustained visual working memory of the object when another person could still see it.

The finding that this visual response to the object was sustained and, in infants, even reactivated after full object occlusion suggests that, when seen by an agent, objects are kept online, or reactivated, in visual working memory, where they are processed similar to perception. This is supported by the specificity of this effect to 4 Hz, which, as a spontaneous intrinsic rhythm in the brain, supports working memory (*64–66*). It is important to note that this effect cannot reflect a general increase in working memory load, as such an intrinsic cognitive process would have led to an increase in *induced* 4 Hz oscillations at arbitrary phases. In contrast, here we observed an increase in *evoked* 4 Hz signal, that is, *phase-locked* 4 Hz activity at synchronized phases. Such phase-locked responses do not occur spontaneously without rhythmic visual stimulation but are a direct perceptual response to a flickering stimulus (*26*, *30*).

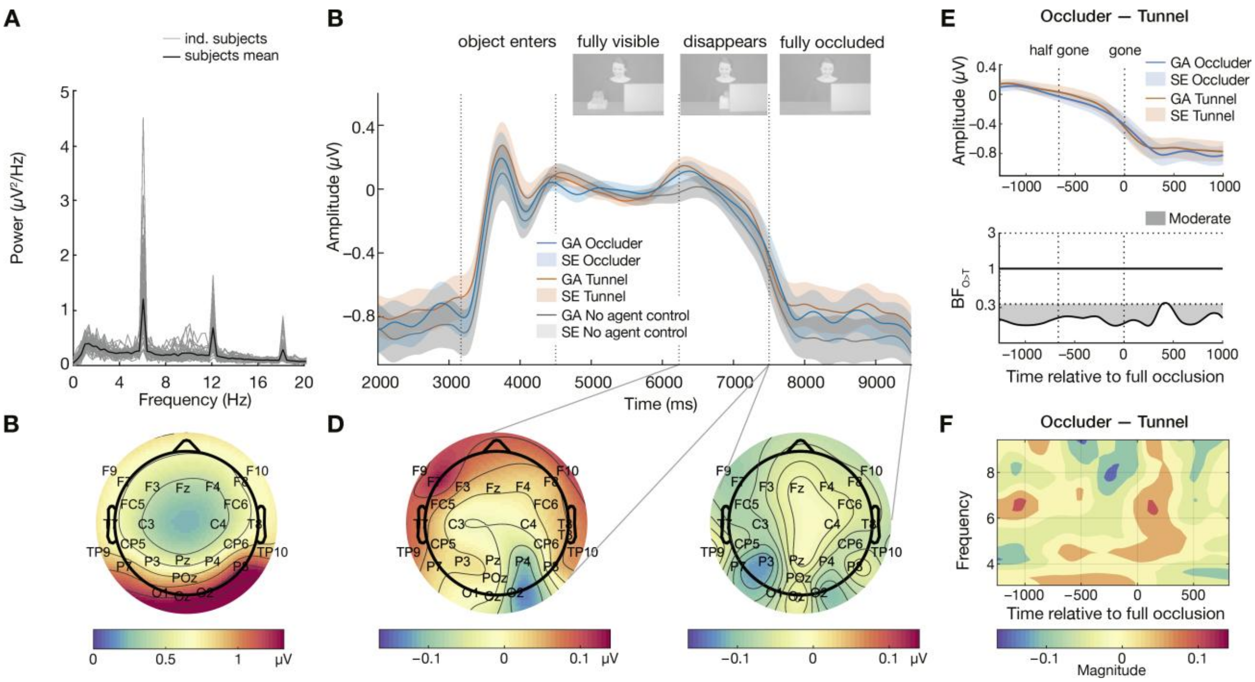


**Fig. S4. Results of evoked oscillations at 6 Hz in *N* = 40 adults.** (**A**) The frequency spectrum shows clear peaks at 6 Hz and its harmonics (electrode Oz) while the object is visible to the participant. (**B**) The topographical results demonstrate a clear 6 Hz response to the visible flickering object over visual cortex. (**C**) Baseline-corrected average 6 Hz response over occipital sensors (O1, O2, Oz) for the Occluder (blue) vs Tunnel condition (orange). The 6 Hz amplitude envelope mirrored the object's visibility. In contrast to 4 Hz, there was no difference between the grand averages (GA) in the Tunnel and Occluder condition, nor with the additional No Agent control condition. This indicates that the prolongation of the signal as a function of the agent's visual access was specific to 4 Hz. Shading indicates the standard error (SE). (**D**) Topography of the evoked 6 Hz Occluder-Tunnel difference while the object disappears (left) and during full occlusion (right). (**E**) Evoked 6 Hz amplitude envelope in the time window of 6234-8500 ms (top) and the respective $BF_{10}$ of the dynamical analysis (bottom) with no difference between the Occluder and Tunnel trials ($BF_{10}$ between .17 and .30). (**F**) The time-frequency plot displays the power difference between the Occluder and Tunnel conditions at the end of the trial.

*S7. Experiment 3 & 4: Eye-tracking*

**Table S1.**

| | | *Disappearance phase* | | | *Full occlusion phase* | | |
|---|---|---|---|---|---|---|---|
| | | Occluder | Tunnel | No Agent | Occluder | Tunnel | No Agent |
| | RLS | *M* (*SD*) | *M* (*SD*) | *M* (*SD*) | *M* (*SD*) | *M* (*SD*) | *M* (*SD*) |
| Adults | $RLS_{face>obj}$ | .12 (.15) | .12 (.16) | .08 (.21) | .61 (.29) | .58 (.32) | .18 (.31) |
| | $RLS_{face>occl/tunnel}$ | .88 (.23) | .85 (.29) | .48 (.45) | .82 (.22) | .83 (.28) | .59 (.37) |
| | $RLS_{obj>occl/tunnel}$ | .99 (.03) | .99 (.03) | .94 (.12) | .68 (.33) | .83 (.18) | .85 (.28) |
| Infants | $RLS_{face>obj}$ | .30 (.36) | .21 (.33) | .20 (.40) | .75 (.25) | .65 (.26) | .41 (.30) |
| | $RLS_{face>occl/tunnel}$ | .89 (.28) | .80 (.35) | .51 (.51) | .91 (.16) | .90 (.16) | .59 (.34) |
| | $RLS_{obj>occl/tunnel}$ | .96 (.13) | .92 (.19) | .94 (.10) | .69 (.37) | .77 (.32) | .66 (.38) |

**Table S1. Descriptive results of Experiment 3 and 4.** The mean and standard deviation of relative looking scores (RLS) across the three AOIs by condition for $N = 24$ adults and $N = 22$ infants averaged across the time window during the disappearance phase (6234-7500 ms) and the full occlusion phase (7500-9500 ms). Note that comparison's involving the face AOI in the no agent condition trivially differ from the other conditions as no face was visible in this condition.

**Table S2.**

| | | Phase | Occluder vs Tunnel ($BF_{10}$) | Occluder vs No Agent ($BF_{10}$) | Tunnel vs No Agent ($BF_{10}$) |
|---|---|---|---|---|---|
| Adults | $RLS_{face>obj}$ | Disappearance | 1.18 | 2.05 | 2.77 |
| | | Full occlusion | .49 | 1780.75 | 1442.61 |
| | $RLS_{face>occl/tunnel}$ | Disappearance | .55 | 22.03 | 75.34 |
| | | Full occlusion | .09 | 22.37 | 39.28 |
| | $RLS_{obj>occl/tunnel}$ | Disappearance | .24 | 1.05 | 1.12 |
| | | Full occlusion | **4.24** | **25.98** | 0.32 |
| Infants | $RLS_{face>obj}$ | Disappearance | **5.12** | 615.69 | 95.07 |
| | | Full occlusion | **5.02** | 217.28 | 88.85 |
| | $RLS_{face>occl/tunnel}$ | Disappearance | .51 | 6.22 | 3.35 |
| | | Full occlusion | .27 | 38.11 | 61.52 |
| | $RLS_{obj>occl/tunnel}$ | Disappearance | .27 | .41 | .28 |
| | | Full occlusion | .30 | .36 | .79 |

**Table S2. Statistical results of Experiment 3 and 4.** The Bayes Factor ($BF_{10}$) comparing the relative looking scores (RLS) between conditions across the three AOIs for the adult and infant samples. The RLS was averaged over the disappearance phase (6234-7500 ms) and the full occlusion phase (7500-9500 ms). Note that comparison's involving the face AOI in the no agent condition trivially differ from the other conditions as no face was visible in this condition and are therefore indicated in light gray. Non-trivial comparisons with evidence for an effect ($BF_{10} > 3$) are marked in bold.

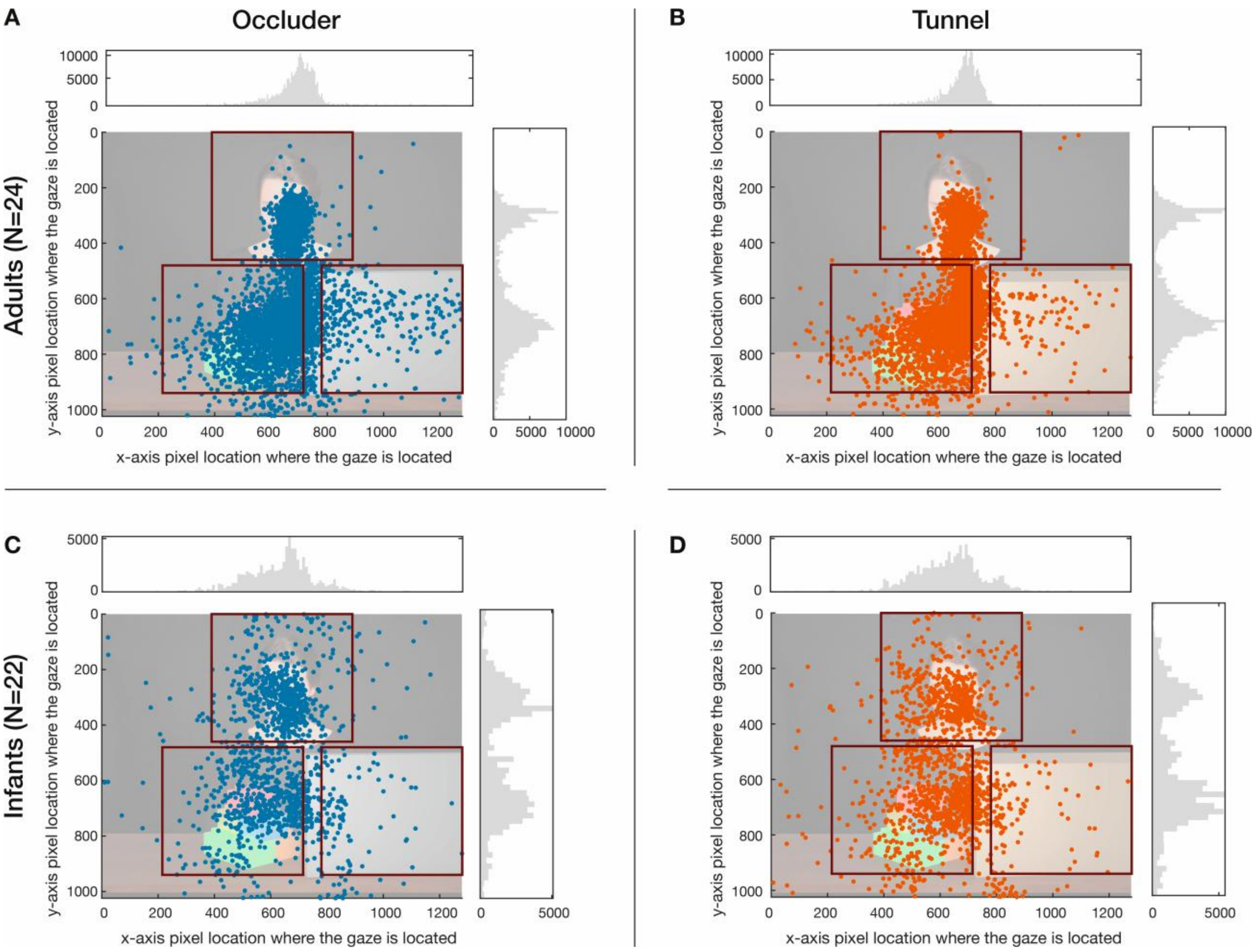


**Fig. S5. Mapped fixation points across all participants and trials of Experiment 3 and 4.** Mapped fixation points and their distributions (gray, summed across all participants and trials) are shown separately for the adult sample (panels **A, B**) and the infant sample (panels **C, D**), for the Occluder condition (**A, C**, blue dots) and Tunnel condition (**B, D**, orange dots) in a time window starting from 4500 ms until the end of the trial and areas of interest (AOI).

*S8. Bayesian sequential testing scheme*

In adults, our Bayesian sequential testing scheme converged towards our predefined stopping criterion after $N$=31 participants. Specifically, the evoked 4 Hz signal converged towards moderate evidence for a difference between the Occluder vs Tunnel conditions in the disappearance phase ($BF_{10}$ = 3.18, W = 345, $\hat{R}$ = 1.003, see Fig. S6A). The sequential Bayes factors from our predefined minimum N to the full sample at $N$ = 40 are shown in Fig. S6B. The results at the stopping criterion of $N$ = 31 were highly similar to the data with the full sample.

In infants, the analyses reached the predefined stopping criterion at $N$ = 40. Specifically, the evoked 4 Hz response converged towards moderate evidence against a difference between the Tunnel and Occluder condition in the averaged disappearance phase ($BF_{10}$ = .15, W = 280, $\hat{R}$ = 1.000), yielding highly similar results to the full sample (see Fig. S6C). Figure S6D shows the sequential Bayes factor from the predefined minimum $N$ of 40 to the full sample.

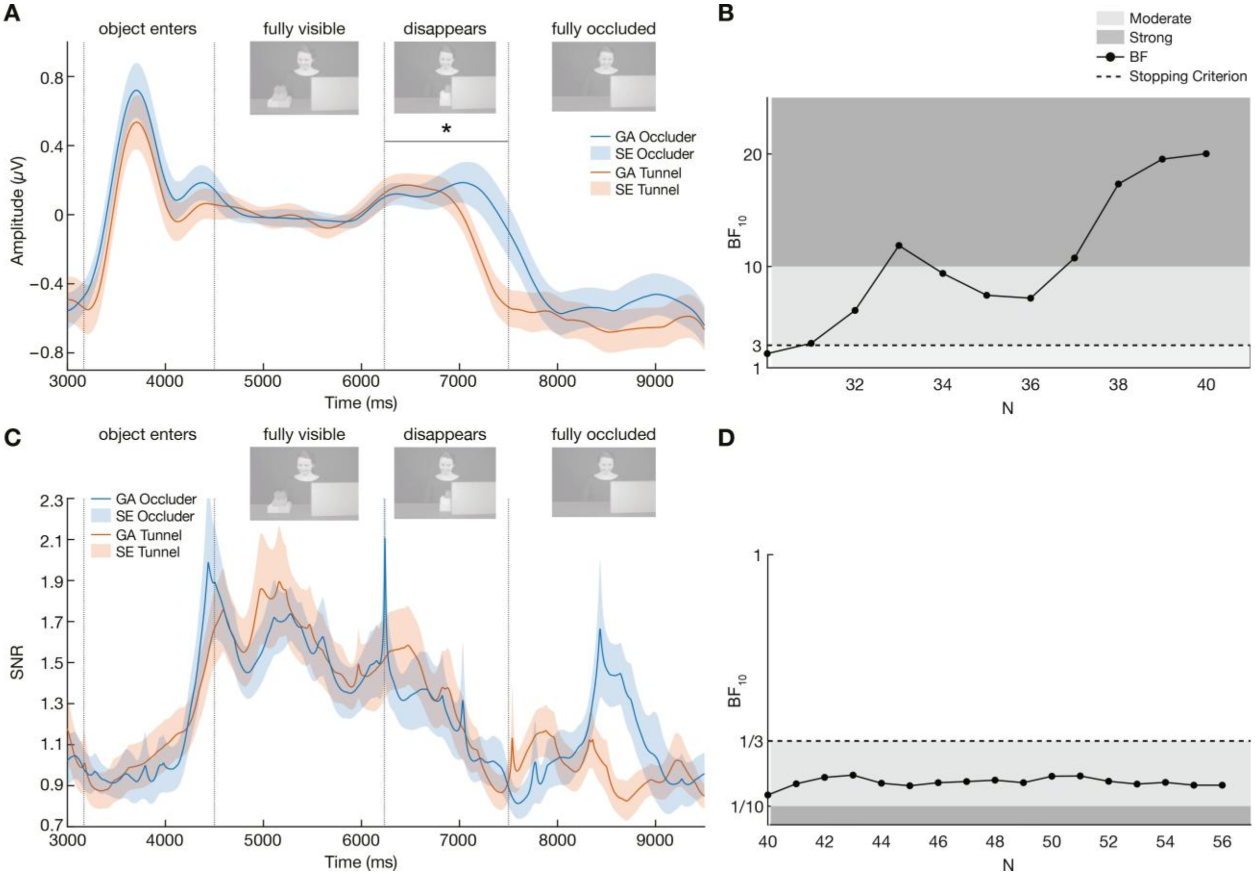


**Fig. S6. Results of adult and infant stop sample according to Bayesian sequential testing**. (**A**) Baseline-corrected grand average (GA) time course of the 4 Hz signal envelope by condition for $N = 31$ adults, averaged over occipital electrodes (Oz, O1, O2). Shading indicates the standard error (SE). (**B**) Sequential Bayes Factor $BF_{10}$ for the Bayesian Wilcoxon Signed Rank test comparing the Tunnel and Occluder condition, averaged over the time window of object disappearance, from $N = 30$ to $N = 40$ (final sample size). The predefined threshold is indicated by the dashed line ($BF_{10} = 3$ and $BF_{10} = 1/3$). (**C**) Signal-to-Noise-Ratio (SNR) of the 4 Hz signal by condition for $N = 40$ infants, averaged over occipital electrodes (Oz, O1, O2). Shading indicates the standard error. (**D**) Resulting Bayes Factor $BF_{10}$ for the Bayesian Wilcoxon Signed Rank test comparing the Occluder vs Tunnel response (stopping criterion) during the disappearance phase from $N = 40$ to $N = 56$ (final sample size). The predefined threshold is indicated by the dashed line ($BF_{10} = 3$ and $BF_{10} = 1/3$).

**Table S3.**

| | Experiment | *N* | female | male | divers | Age (*Median* (*Range*)) | Trial number Occluder *M* (*SD*) | Trial number Tunnel *M* (*SD*) | Trial number No Agent *M* (*SD*) |
|---|---|---|---|---|---|---|---|---|---|
| EEG | Exp 1: Adults | 40 | 15 | 25 | 0 | *Median* = 26 years, *range*: 19 - 34 | 28.6 (± 1.6) | 28.2 (± 1.9) | 28.5 (± 2) |
| | Exp 2: Infants | 56 | 31 | 25 | 0 | *Median* = 13.4 months, *range* = 12.1 – 15.4 | 11.4 (± 6.1) | 12 (± 8.2) | 8.6 (± 7.4) |
| Eye-tracking | Exp 3: Adults | 24 | 13 | 9 | 2 | *Median* = 25.5 years, *range* = 19 – 36 | 29.8 (± .2) | 29.8 (± .3) | 29.8 (± .2) |
| | Exp 4: Infants | 22 | 11 | 11 | 0 | *Median* = 13.8 months, *range* = 12.1 – 15.5 | 19.4 (± .6) | 19.8 (± .5) | 19.7 (± .5) |

**Table S3. Descriptive statistics for the four experiments**. Overview of sample size, demographics, and trial count per condition for all four experiments.

*S9. Distribution plots*

As q-q-plots indicated minor deviations from normality for both the adult and infant sample (see Fig. S7), we used non-parametric statistical test alternatives whenever possible.

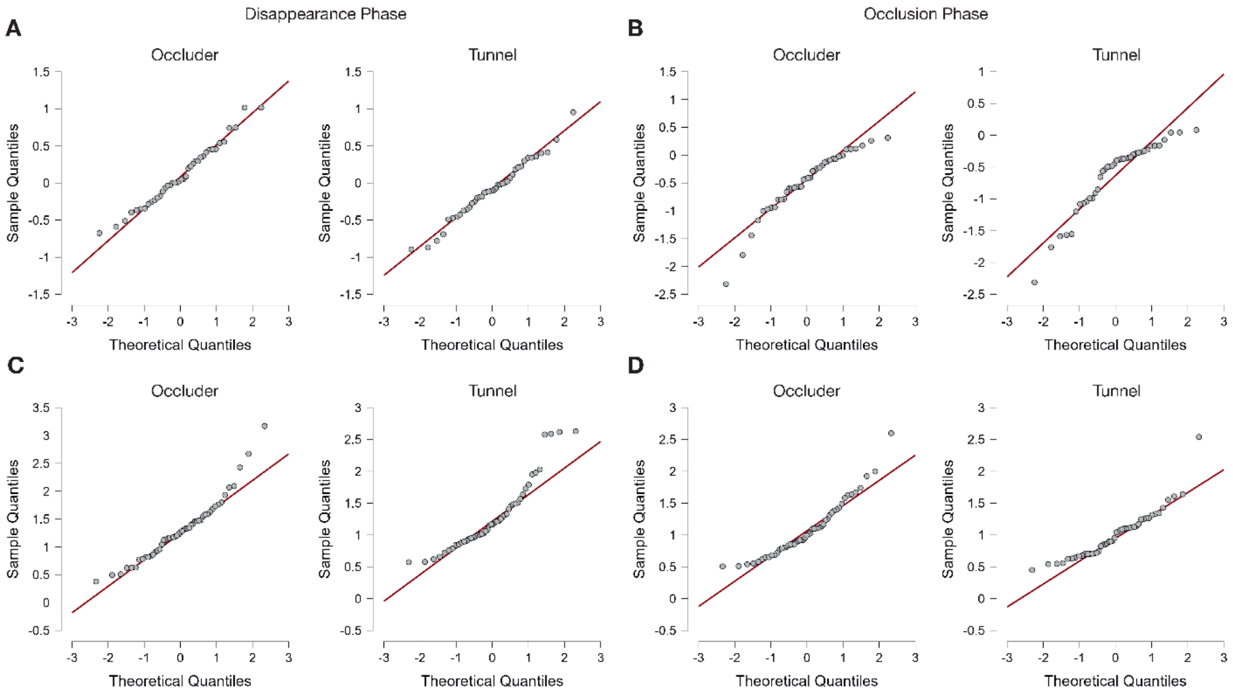


**Fig. S7. Distribution plots for the averaged 4 Hz response.** Distribution plots for the Occluder and Tunnel condition averaged over the disappearance phase and full occlusion phase for $N = 40$ adults (**A**, **B**) and $N = 56$ infants (**C**, **D**).

*S10. Looking duration - Experiment 2*

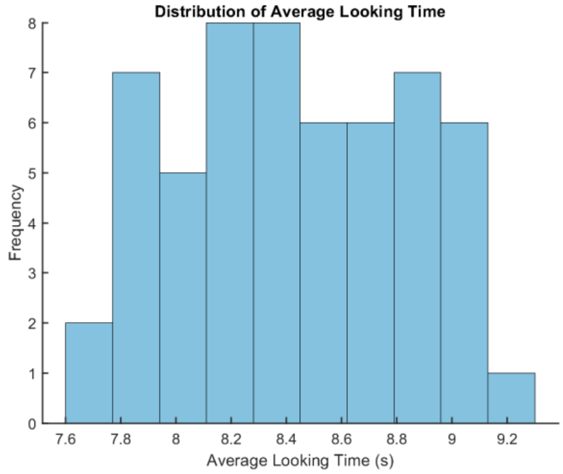


**Fig. S8. Distribution of the average looking duration per trial for Experiment 2 (Infant EEG).** The average looking duration across infants for included trials was *M* = 8.43 sec, *SD* = 0.42.

**Table S4.**

| Minimum looking duration per trial | 6.5 sec | 7.5 sec | 8.5 sec |
|---|---|---|---|
| Phase | Full object phase and half of the disappearance phase | Full object phase and full disappearance phase | Full object phase, full object disappearance phase, and half of the full occlusion phase |
| Time window | 8524 - 8636 ms | 8400 - 8652 ms | 8556 - 8688 ms |
| $BF_{10}$ | 3 - 6.35 | 3 - 7.54 | 3 - 11.34 |

**Table S4. Looking duration Experiment 2 - Comparison of different inclusion criteria.** The time window and Bayes Factor ($BF_{10}$) indicating a greater response in the Occluder compared to the Tunnel condition across three different coding criteria for the infant sample of Experiment 2. The effect time window showed substantial overlap using different trial inclusion criteria.

**Movie S1.**
Exemplary video of Occluder condition. Objects flickering at 4 Hz disappeared behind an occluder, providing continued visual access for the agent.

**Movie S2.**
Exemplary video of Tunnel control condition. Objects flickering at 4 Hz disappeared inside a tunnel, so that neither participant nor agent had visual access to the object.

**Movie S3.**
Exemplary video of additional No Agent control condition. This control was identical to the Occluder condition, except that a blurred light was shown instead of an agent that matched the luminance of the agent conditions.

**Movie S4.**
Exemplary video of the Familiarization video for the Occluder condition. The Occluder entered the scene, made an 180° turn when it reached the middle of the scene, and then moved into its final position, while the agent followed it with her gaze.

**Movie S5.**
Exemplary video of the Familiarization video for the Tunnel condition. The Tunnel entered the scene, made an 180° turn when it reached the middle of the scene, and then moved into its final position, while the agent followed it with her gaze.
